# Changes in presynaptic gene expression during homeostatic compensation at a central synapse

**DOI:** 10.1101/2020.11.25.398370

**Authors:** Evan R. Harrell, Diogo Pimentel, Gero Miesenböck

**Author notes:** **Declaration of Interests:** The authors declare no competing financial interests.

## Abstract

Homeostatic matching of pre- and postsynaptic function has been observed in many species and neural structures, but whether transcriptional changes contribute to this form of trans-synaptic coordination remains unknown. To identify genes whose expression is altered in presynaptic neurons as a result of perturbing postsynaptic excitability, we applied a transcriptomics-friendly, temperature-inducible Kir2.1-based activity clamp at the first synaptic relay of the *Drosophila* olfactory system, a central synapse known to exhibit trans-synaptic homeostatic matching. Twelve hours after adult-onset suppression of activity in postsynaptic antennal lobe projection neurons, we detected changes in the expression of many genes in the third antennal segment, which houses the somata of presynaptic olfactory receptor neurons. These changes affected genes with roles in synaptic vesicle release and synaptic remodeling, including several genes implicated in homeostatic plasticity at the neuromuscular junction. At 48 hours and beyond, the transcriptional landscape was tilted toward proteostasis, energy metabolism, and cellular stress defenses, indicating that the system had been pushed to its homeostatic limits. Our data provide insights into the nature of homeostatic compensation at a central synapse and identify many genes engaged in synaptic homeostasis. The presynaptic transcriptional response to genetically targeted postsynaptic perturbations could be exploited for the construction of novel connectivity tracing tools.

**Significance Statement:** Homeostatic feedback mechanisms adjust intrinsic and synaptic properties of neurons to keep their average activity levels constant. We show that, at a central synapse in the fruit fly brain, these mechanisms include changes in presynaptic gene expression that are instructed by an abrupt loss of postsynaptic excitability. The trans-synaptically regulated genes have roles in synaptic vesicle release and synapse remodeling; protein synthesis, folding, and degradation; and energy metabolism. Our analysis suggests that similar homeostatic machinery operates at peripheral and central synapses, identifies some of its components, and potentially opens new opportunities for the development of connectivity-based gene expression systems.

## Introduction

Homeostatic feedback that stabilizes network activity after synaptic weight changes is an important adjunct to correlation-based learning rules (Turrigiano, 2011). Early demonstrations of homeostatic plasticity followed pharmacological manipulations of synaptic transmission in neuronal cultures (Turrigiano et al., 1994, 1998). When global activity levels were artificially increased or decreased, homeostatic forces intervened to maintain firing rates within defined ranges. These homeostatic forces are generated by two processes (Turrigiano, 2011): cell-autonomous changes in intrinsic excitability, which alter the gain of the neuronal voltage response to synaptic currents (Turrigiano et al., 1994; Desai et al., 1999); and adjustments of the synaptic strengths themselves (Petersen et al., 1997; Davis et al., 1998; Turrigiano et al., 1998; Burrone et al., 2002). These adjustments, though in principle also achievable in a cell-autonomous fashion by altering the density of neurotransmitter receptors in the postsynaptic membrane (Wierenga et al., 2005; Goold and Nicoll, 2010), often involve a trans-synaptic partnership in which postsynaptic neurons communicate deviations from their activity setpoint via retrograde signals to their presynaptic partners, which in turn increase or decrease transmitter release (Cull-Candy et al., 1980; Petersen et al., 1997; Sandrock et al., 1997; Davis et al., 1998; Burrone et al., 2002; Haghighi et al., 2003; Thiagarajan et al., 2005).

Much existing knowledge of retrograde communication comes from studies of the neuromuscular junction (NMJ). In mammals and *Drosophila*, mutations or autoantibodies that reduce the responsiveness of muscle to neurotransmitter cause compensatory increases in motor neuron vesicular release (Cull-Candy et al., 1980; Petersen et al., 1997; Sandrock et al., 1997; Davis et al., 1998). At the *Drosophila* NMJ, acute pharmacological receptor blockade (Frank et al., 2006) or expression of the inwardly rectifying potassium channel Kir2.1 in muscle (Paradis et al., 2001) induce similar presynaptic compensatory effects. While many gene products and signaling pathways have been implicated in synaptic homeostasis (Davis and Müller, 2015), knowledge of the transcriptional changes that may be required to lock the presynaptic cells into their altered functional state remains scant (Marie et al., 2010).

Pre- and postsynaptic function are also matched at the central synapses between olfactory receptor neurons (ORNs) and projection neurons (PNs) in the antennal lobe of *Drosophila* (Kazama and Wilson, 2008), where the axons of 20–200 ORNs expressing the same odorant receptor connect to dendrites of an average of three affine PNs in a precise anatomical register (Groschner and Miesenböck, 2019). There is clear covariation between the dendritic arbor sizes of PNs belonging to different transmission channels and the amplitudes of unitary excitatory postsynaptic currents (EPSCs): the larger unitary EPSCs of PNs with larger dendritic trees—and, therefore, lower impedances—reflect homeostatic increases in the number of presynaptic ORN release sites in response to increased postsynaptic demand for synaptic drive (Kazama and Wilson, 2008; Mosca and Luo, 2014). This central model of synaptic homeostasis has been characterized physiologically and anatomically, but the molecular mechanisms of synaptic matching are unexplored. Taking advantage of the ease with which the presynaptic partners at this synapse can be isolated (they reside in an external appendage, the third antennal segment), we carried out a transcriptome-wide screen for genes regulated by retrograde homeostatic signals. Homeostatic plasticity was induced by adult-onset expression of Kir2.1 in PNs; the expression of a non-conducting mutant of Kir2.1 (Kir2.1-nc) served as control.

## Methods

### Drosophila strains and culture

Flies were maintained at 21°C and 65% humidity on a constant 12:12-hour light:dark (LD) cycle in rich cornmeal and molasses-based food with brewer’s yeast. Driver lines *GH146-GAL4* (Stocker et al., 1997) and *pdf-GAL4* (Renn et al., 1999) were used to target the expression of codon-optimized *UAS-Kir2.1* transgenes (see below) to PNs and PDF-expressing clock neurons, respectively. Three copies of two *tubulin-GAL80*^*ts*^ insertions on different chromosomes (McGuire et al., 2003) were combined to achieve tight repression of the GAL4-responsive transgenes until induction. The induction incubator was kept at 31°C in 70% humidity on the same 12:12 LD schedule.

The cDNA sequence encoding human Kir2.1 was codon-optimized for *Drosophila* (GenBank accession number MW088713), synthesized at MWG Eurofins, and fused to a codon-optimized N-terminal EGFP tag. The non-conducting variant (Kir2.1-nc) was created by mutating codon 146 of the ion channel sequence from glycine to serine (GGA to AGC) (Haruna et al., 2007). The channel constructs replaced the mCD8::GFP coding sequence in derivatives of plasmid *pJFRC2-10XUAS-IVS-mCD8::GFP* (Pfeiffer et al., 2010), which were inserted into the *attp2* landing site on the third autosome.

### Confocal microscopy

Female flies aged 5 days were anesthetized on ice and dissected in phosphate-buffered saline (PBS; 1.86 mM NaH_2_PO_4_, 8.41 mM Na_2_HPO_4_, 175 mM NaCl). Immediately after dissection, brains were fixed in ice-cold PBS containing 4% (w/v) paraformaldehyde for 1–2 hours at room temperature, rinsed three times in ice-cold PBS containing 0.1% (w/v) Triton X-100 (PBT), washed three times for 20 minutes in ice-cold PBT, and mounted and cleared in Vectashield (Vector Labs). Confocal image stacks with an axial spacing of 1–1.5 μm were collected on a Leica TCS SP5 microscope with an HCX IRAPO L 25x/0.95 W objective.

### Electrophysiology

Targeted whole-cell patch-clamp recordings from the fluorescent somata of PNs expressing EGFP::Kir2.1 or EGFP::Kir2.1-nc were obtained through a small cranial window in 5-day old females. The brain was continuously superfused with extracellular solution containing 103 mM NaCl, 3 mM KCl, 5 mM TES, 8 mM trehalose, 10 mM glucose, 7 mM sucrose, 26 mM NaHCO_3_, 1 mM NaH_2_PO_4_, 1.5 mM CaCl_2_, 4 mM MgCl_2_ (pH 7.3) and equilibrated with 95% O_2_–5% CO_2_. Borosilicate glass electrodes (7–13 MΩ) were filled with intracellular solution containing 140 mM potassium aspartate, 10 mM HEPES, 1 mM KCl, 4 mM Mg-ATP, 0.5 mM Na3GTP, 1 mM EGTA (pH 7.3). Signals were acquired with a MultiClamp 700B Microelectrode Amplifier, filtered at 6–10 kHz, and digitized at 10–20 kHz with an ITC-18 data acquisition board controlled by the Nclamp and NeuroMatic packages. Data were analyzed with NeuroMatic (http://neuromatic.thinkrandom.com) and custom procedures in Igor Pro (WaveMetrics) (Donlea et al., 2014). The membrane time constant was determined by fitting a single exponential to the voltage deflection caused by a 200-ms-long hyperpolarizing current pulse. Input resistances were estimated from linear fits of the subthreshold voltage deflections elicited by small current pulses of increasing amplitude and a duration of 1 s. Firing rates were quantified by holding cells at resting potentials of –60 ± 2 mV and injecting sequences of depolarizing current pulses (5 pA increments, 1 s duration). Spikes were detected by finding minima in the second derivative of the membrane potential record. The spike rate was calculated by dividing the number of action potentials discharged by the time elapsed between the first and last spike. The current amplitude at which each cell reached a given frequency threshold (1–50 Hz) was used to construct cumulative distribution functions. The distributions were fit with logistic Naka-Rushton functions of the form (Donlea et al., 2014):

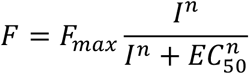

where *F* is the percentage of cells reaching threshold at a given current level *I*, *F*_*max*_ is the percentage of cells reaching threshold at maximal current, *EC*_*50*_ indicates the half-maximal or semisaturation current, and the exponent *n* determines the steepness of the curve. With only two free parameters (*EC*_*50*_ and *n*, given that *F*_*max*_ is measured experimentally), this simple model provided a satisfying fit to all distributions.

### Analysis of circadian behavior

Three-day old female flies were individually inserted into 65 mm glass tubes and loaded into the Trikinetics Drosophila Activity Monitor system, which was operated at 31°C in 24-hour dark (DD) conditions for 5–7 days. Group sizes for activity measurements (16 experimental and 16 control flies) reflect the capacity of the monitors.

### Third antennal segment dissection

Groups of 20–30 flies were aged in precisely controlled temperature conditions for 5 days (see Fig. 4*A*) and decapitated with a surgical scalpel on a CO_2_ pad; the heads were transferred to petri dishes kept on dry ice. Once a petri dish contained ~50 heads, it was sealed with parafilm and stored at −80°C until RNA extraction. The sealed petri dishes were dipped in liquid nitrogen for 60 seconds, vortexed at full strength for 60 seconds, and then unsealed and placed on a dry-ice-chilled glass stand under a dissection microscope. Individual third antennal segments were picked with fine forceps and placed directly into 100 μl TRIzol (Thermo Fisher Scientific).

### RNA extraction

Third antennal segments in 100 μl TRIzol were disrupted with several strokes in a Dounce homogenizer. The homogenates were diluted with 900 μl TRIzol and incubated at room temperature for 5 minutes. Samples destined for 3’ digital gene expression profiling (3’ DGE) underwent phase separation after the addition of 225 μl chloroform; RNA in the aqueous phase was precipitated with isopropanol and resuspended in 5 μl RNase-free water. Total RNA for RNA-seq and RT-qPCR was isolated with the help of RNeasy minelute columns (Qiagen), following the addition of 400 μl of 70% RNase-free ethanol to the TRIzol homogenates and on-column DNaseI digests. Samples were snap frozen in liquid nitrogen and stored at −80°C.

### cDNA library generation

Libraries for 3’ DGE were generated at MWG Eurofins Genomics from ultrasonically fragmented poly(A)-tailed RNA, which was isolated using oligo(dT) chromatography. Following ligation of an RNA adapter to the 5’-end, the mRNA fragments were reverse-transcribed from an oligo(dT) primer, and the resulting cDNA was PCR-amplified with a high-fidelity polymerase. Each cDNA library was purified, size selected, quality-checked by capillary electrophoresis, and sequenced on the HiSeq2000 platform (Illumina) in 1×100 bp run mode.

For RNA-seq and RT-qPCR, oligo(dT)-enriched RNA underwent 14 cycles of amplification using the SMARTer Ultra Low RNA Kit for Illumina Sequencing (Clontech). After cDNA fragmentation, libraries were prepared in an additional 15 amplification cycles using the NEBNext Ultra DNA Library Prep Kit for Illumina (New England Biolabs) and sequenced on the HiSeq2000 platform (Illumina) in paired-end mode.

### Transcriptome analysis

Raw reads were 100 base pairs (bp) in length (paired-end reads for RNA-seq and single-ended reads for 3’ DGE). Fastq files, containing reads and quality scores, were first run through the FastQC package (Andrews, 2010). Highly abundant sequences that did not map to the *Drosophila* genome (and originated from primers or amplification artefacts) were eliminated using Trimmomatic software (Bolger et al., 2014). Reads were scanned with a 4-bp sliding window and cut when the average quality dropped below 15; trimmed reads shorter than 25 bp were discarded. The reads were mapped to Ensembl DM genome release 5.74 using TopHat2 (Kim et al., 2013), assigned to transcripts annotated in the transcript file of the Berkeley Drosophila Genome Project (BDGP) release 5.74 with Cufflinks, and merged into an experiment-wide gtf file with Cuffmerge (Trapnell et al., 2012). The gtf file was used to produce raw read counts (using HTSeq) suitable for differential expression analysis in DESeq2 (Love et al., 2014). The topGO and *ViSEAGO* packages were used to analyze the enrichment of gene ontology (GO) terms in the set of differentially expressed genes called by DESeq2 (unadjusted *p* < 0.05) vis-à-vis a reference set of all genes with a normalized expression level above 1 (the “gene universe”) (Alexa et al., 2006; Brionne et al., 2019). To keep the number of Fisher’s exact tests to a minimum, only GO terms with more than 40 attached genes were considered. Enriched GO terms with unadjusted *p* < 0.01 were clustered hierarchically according to Wang's distance, a measure of semantic similarity (Wang et al., 2007; Brionne et al., 2019).

### Real time quantitative PCR (RT-qPCR)

Transcript levels were determined by quantitative real-time PCR on a LightCycler 480 system (Roche) using SYBR Green I Master Mix (Roche) in 10-μl reactions containing 100 nM of each gene-specific primer and 50 ng of pre-amplified cDNA. Two sets of primers were designed for each gene of interest. All samples were run in technical triplicates; non-reverse-transcribed mRNA and water served as negative controls. Melting curves were analyzed after amplification, and amplicons were visualized by agarose gel electrophoresis to confirm primer specificity. Relative transcript levels were estimated with the help of the 2^−ΔΔ*C*t^ method (Livak and Schmittgen, 2001), using the housekeeping gene *CycK* for normalization.

## Results

### Antennal transcriptomics

To characterize gene expression in the third antennal segment, 5-day old male Canton-S (CS) flies were decapitated either between zeitgeber time (ZT) 5 and ZT8 (the day group) or between ZT17 and ZT20 (the night group). After snap-freezing, third antennal segments were manually isolated, and total RNA was extracted in a single batch to minimize variability (Fig. 1*A*, see Methods). For both day and night conditions, three biological replicates were prepared, and the resulting six cDNA libraries were sequenced on one lane of an Illumina HiSeq2000 machine using 3’ digital gene expression profiling (3’ DGE) technology. After stringent quality assessment and read trimming (Fig. 1*A*), the high-quality reads were mapped to the *Drosophila* genome (for mapping statistics, see Table 1). Biological replicates showed high correlations with one another (Fig. 1*B*, Table 1), and day and night samples could easily be distinguished on the basis of their top two principal components (Fig. 1*B*, inset). Underlying this clean separability were 128 differentially expressed genes, identified by DESeq2 (Love et al., 2014) with a false discovery rate (FDR)-adjusted significance level of < 0.2, and large expression level differences between the day and night (Fig. 1*C*, Table 2). Core clock components, such as *cryptochrome*, *Clock*, *period*, *timeless,* and *vrille*, were found near the top of the amplitude distribution of oscillating transcripts (Fig. 1*C*), in two groups at opposite poles of the 24-hour cycle, consistent with their antagonistic roles in the transcriptional feedback oscillator (Claridge-Chang et al., 2001; McDonald and Rosbash, 2001).

**Figure 1.**
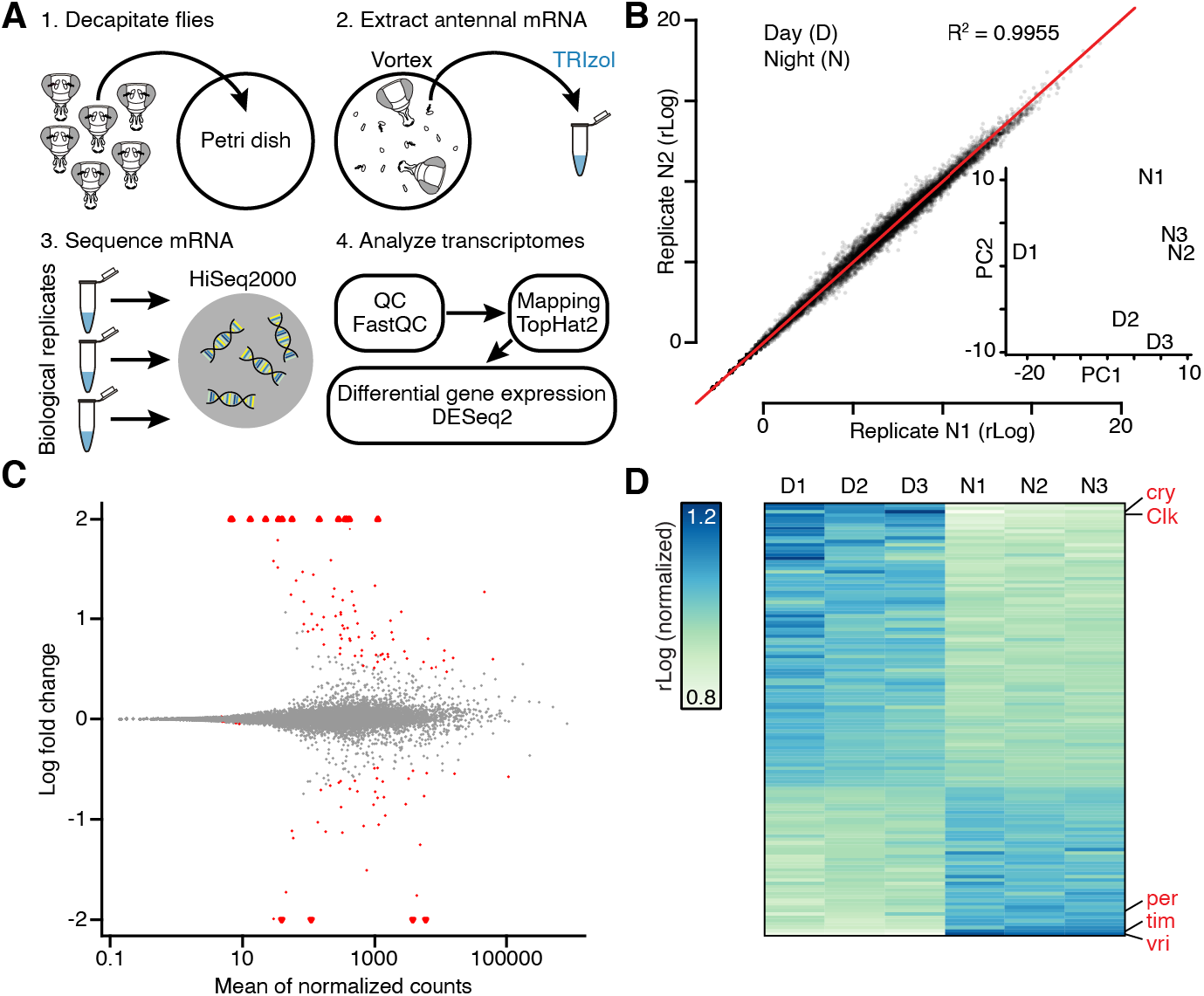
Antennal transcriptomics: workflow, diagnostics, and functional validation. ***A***, Experimental workflow. ***B***, Scatterplot of gene expression levels in biological replicates N1 vs. N2. Inset: D and N samples in a principal component analysis (PCA) plot. ***C***, MA plot of log_2_ fold change in expression vs. mean expression level of all transcripts. Triangles represent data points outside the plotted range. ***D***, Expression levels of all transcripts with FDR-adjusted *p* < 0.20 during the day and night. Each column represents a sequencing library generated from third antennal segments. Core clock components are indicated in red.

**Table 1.** Mapping metrics of third antennal segment transcriptomes collected during the day (D) and night (N)

| ID | *n* Reads | *n* Mapped | % Mapped | *n* Unique | % Unique | Average R<sup>2</sup> |
| --- | --- | --- | --- | --- | --- | --- |
| D1 | 20,857,542 | 14,732,179 | 70.63% | 13,342,850 | 63.97% | 0.9897 |
| D2 | 19,466,527 | 14,408,098 | 74.01% | 13,373,355 | 68.70% | 0.9931 |
| D3 | 18,908,595 | 14,656,324 | 77.51% | 13,440,037 | 71.08% | 0.9919 |
| N1 | 22,905,480 | 18,406,329 | 80.36% | 17,187,773 | 75.04% | 0.9955 |
| N2 | 21,095,058 | 16,864,414 | 79.94% | 15,564,220 | 73.78% | 0.9963 |
| N3 | 23,547,296 | 18,429,070 | 78.26% | 16,918,247 | 71.85% | 0.9963 |
| Average | 21,130,083 | 16,249,402 | 76.79% | 14,971,080 | 70.74% | 0.9938 |

Absolute number and percentages of reads aligned to the transcript file of the Berkeley Drosophila Genome Project (BDGP) release 5.74. *n* Reads: number of raw reads; *n* Mapped and % Mapped: number of reads mapping to the reference transcriptome with corresponding percentage; *n* Unique and % Unique: number of reads mapping to a unique location in the reference transcriptome with corresponding percentage; Average R<sup>2</sup>: average replicate correlation (Pearson's correlation coefficient calculated across all genes with a non-zero read count in at least one replicate).

**Table 2.**
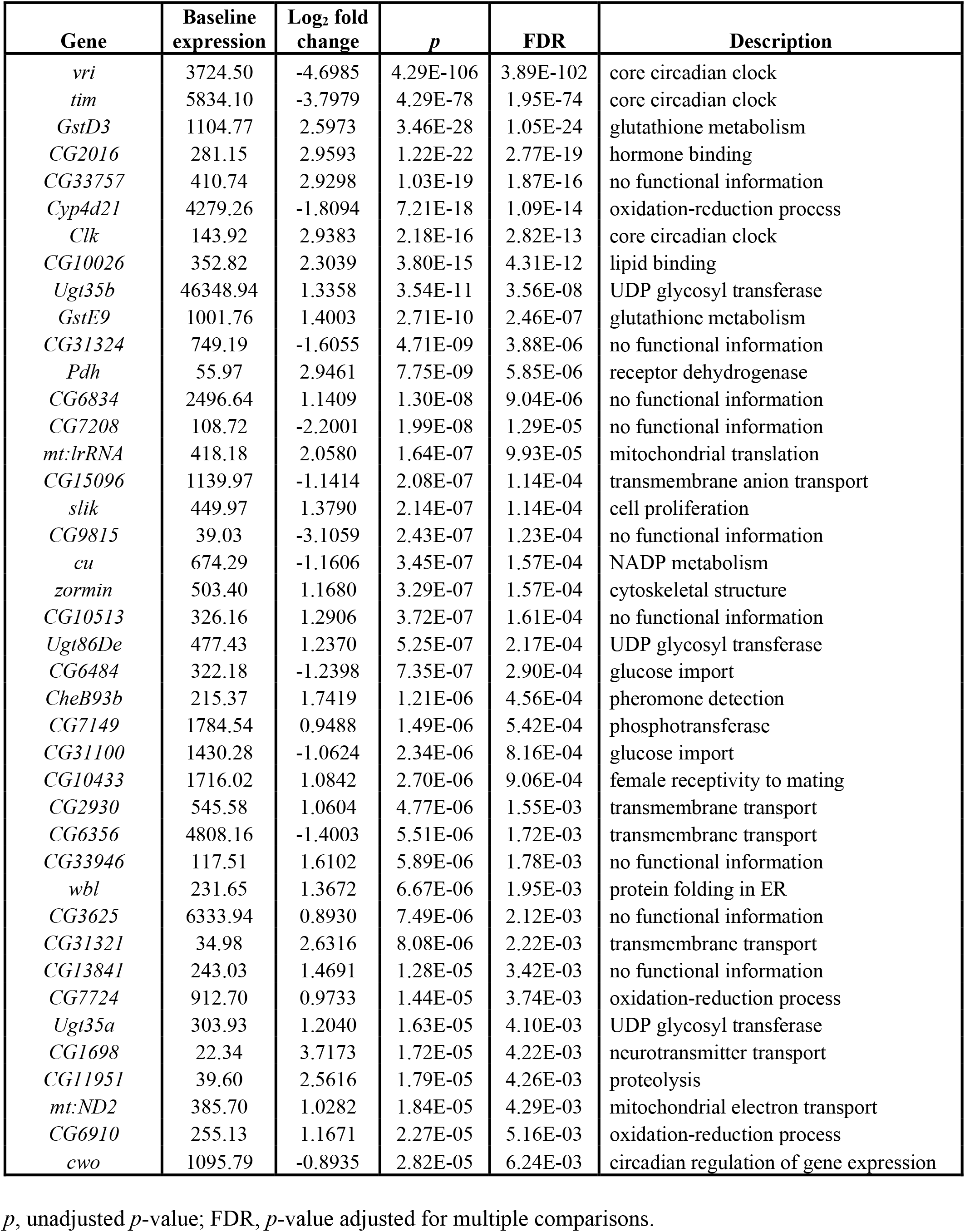
Circadian-regulated genes (FDR-adjusted p < 0.01)

| Gene | Baseline expression | Log<sub>2</sub> fold change | $p$ | FDR | Description |
| --- | --- | --- | --- | --- | --- |
| *vri* | 3724.50 | -4.6985 | 4.29E-106 | 3.89E-102 | core circadian clock |
| *tim* | 5834.10 | -3.7979 | 4.29E-78 | 1.95E-74 | core circadian clock |
| *GstD3* | 1104.77 | 2.5973 | 3.46E-28 | 1.05E-24 | glutathione metabolism |
| *CG2016* | 281.15 | 2.9593 | 1.22E-22 | 2.77E-19 | hormone binding |
| *CG33757* | 410.74 | 2.9298 | 1.03E-19 | 1.87E-16 | no functional information |
| *Cyp4d21* | 4279.26 | -1.8094 | 7.21E-18 | 1.09E-14 | oxidation-reduction process |
| *Clk* | 143.92 | 2.9383 | 2.18E-16 | 2.82E-13 | core circadian clock |
| *CG10026* | 352.82 | 2.3039 | 3.80E-15 | 4.31E-12 | lipid binding |
| *Ugt35b* | 46348.94 | 1.3358 | 3.54E-11 | 3.56E-08 | UDP glycosyl transferase |
| *GstE9* | 1001.76 | 1.4003 | 2.71E-10 | 2.46E-07 | glutathione metabolism |
| *CG31324* | 749.19 | -1.6055 | 4.71E-09 | 3.88E-06 | no functional information |
| *Pdh* | 55.97 | 2.9461 | 7.75E-09 | 5.85E-06 | receptor dehydrogenase |
| *CG6834* | 2496.64 | 1.1409 | 1.30E-08 | 9.04E-06 | no functional information |
| *CG7208* | 108.72 | -2.2001 | 1.99E-08 | 1.29E-05 | no functional information |
| *mt:lrRNA* | 418.18 | 2.0580 | 1.64E-07 | 9.93E-05 | mitochondrial translation |
| *CG15096* | 1139.97 | -1.1414 | 2.08E-07 | 1.14E-04 | transmembrane anion transport |
| *slik* | 449.97 | 1.3790 | 2.14E-07 | 1.14E-04 | cell proliferation |
| *CG9815* | 39.03 | -3.1059 | 2.43E-07 | 1.23E-04 | no functional information |
| *cu* | 674.29 | -1.1606 | 3.45E-07 | 1.57E-04 | NADP metabolism |
| *zormin* | 503.40 | 1.1680 | 3.29E-07 | 1.57E-04 | cytoskeletal structure |
| *CG10513* | 326.16 | 1.2906 | 3.72E-07 | 1.61E-04 | no functional information |
| *Ugt86De* | 477.43 | 1.2370 | 5.25E-07 | 2.17E-04 | UDP glycosyl transferase |
| *CG6484* | 322.18 | -1.2398 | 7.35E-07 | 2.90E-04 | glucose import |
| *CheB93b* | 215.37 | 1.7419 | 1.21E-06 | 4.56E-04 | pheromone detection |
| *CG7149* | 1784.54 | 0.9488 | 1.49E-06 | 5.42E-04 | phosphotransferase |
| *CG31100* | 1430.28 | -1.0624 | 2.34E-06 | 8.16E-04 | glucose import |
| *CG10433* | 1716.02 | 1.0842 | 2.70E-06 | 9.06E-04 | female receptivity to mating |
| *CG2930* | 545.58 | 1.0604 | 4.77E-06 | 1.55E-03 | transmembrane transport |
| *CG6356* | 4808.16 | -1.4003 | 5.51E-06 | 1.72E-03 | transmembrane transport |
| *CG33946* | 117.51 | 1.6102 | 5.89E-06 | 1.78E-03 | no functional information |
| *wbl* | 231.65 | 1.3672 | 6.67E-06 | 1.95E-03 | protein folding in ER |
| *CG3625* | 6333.94 | 0.8930 | 7.49E-06 | 2.12E-03 | no functional information |
| *CG31321* | 34.98 | 2.6316 | 8.08E-06 | 2.22E-03 | transmembrane transport |
| *CG13841* | 243.03 | 1.4691 | 1.28E-05 | 3.42E-03 | no functional information |
| *CG7724* | 912.70 | 0.9733 | 1.44E-05 | 3.74E-03 | oxidation-reduction process |
| *Ugt35a* | 303.93 | 1.2040 | 1.63E-05 | 4.10E-03 | UDP glycosyl transferase |
| *CG1698* | 22.34 | 3.7173 | 1.72E-05 | 4.22E-03 | neurotransmitter transport |
| *CG11951* | 39.60 | 2.5616 | 1.79E-05 | 4.26E-03 | proteolysis |
| *mt:ND2* | 385.70 | 1.0282 | 1.84E-05 | 4.29E-03 | mitochondrial electron transport |
| *CG6910* | 255.13 | 1.1671 | 2.27E-05 | 5.16E-03 | oxidation-reduction process |
| *cwo* | 1095.79 | -0.8935 | 2.82E-05 | 6.24E-03 | circadian regulation of gene expression |

$p$ , unadjusted $p$ -value; FDR, $p$ -value adjusted for multiple comparisons.

Transcripts encoding olfactory, gustatory, and ionotropic receptors (ORs, GRs, and IRs) provided an index of the purity of our library preparations. Certain ORs, IRs, and GRs are expressed in antennal ORNs but not elsewhere, while others are absent from antennal ORNs but present in different types of sensory neuron (Clyne et al., 1999; Gao and Chess, 1999; Vosshall et al., 1999; Scott et al., 2001; Benton et al., 2009). We detected the former, but not the latter, members of all three receptor families in abundance (Fig. 2*A*–*C*).

**Figure 2.**
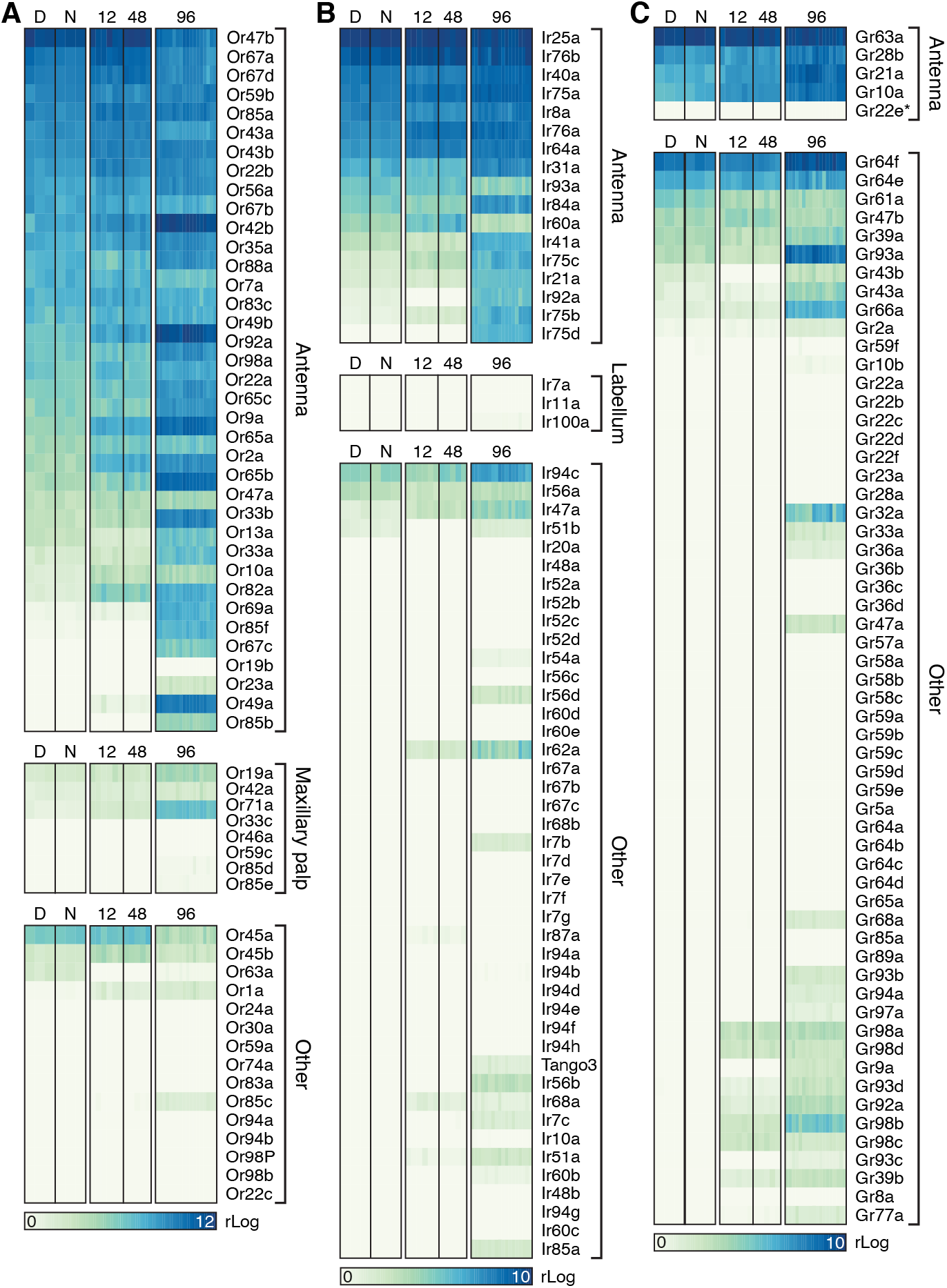
Expression levels of ORs (***A***), IRs (***B***), and GRs (***C***) in sequencing libraries generated from third antennal segments. Each column represents a library generated during the day (D) or night (N), or after 12-hour, 48-hour, or 96-hour induction of Kir2.1 or Kir2.1-nc. The gene encoding the obligatory OR coreceptor Orco/Or83b was expressed at a level above those of other *OR* genes (mean rLog ± SEM = 13.6178 ± 0.16503) and omitted from ***A***.

### Transcriptomics-friendly manipulation of postsynaptic excitability

Kir2.1 is an inwardly rectifying potassium channel that decreases the input resistance of neurons and clamps their membrane potential at or below its resting value; it is widely used as a neuronal “silencer” (Johns et al., 1999). Some single amino acid substitutions in the P-loop signature sequence of the channel (Heginbotham et al., 1994), such as G146S (here called Kir2.1-nc), block ion flow without affecting the protein’s localization (Haruna et al., 2007). We generated *Drosophila* codon-optimized *UAS-EGFP::Kir2.1* and *UAS-EGFP::Kir2.1-nc* lines and crossed them to the *GH146-GAL4* driver, which directs transgene expression to PNs (Stocker et al., 1997). Whereas the non-conducting Kir2.1-nc variant proved innocuous, the expression of functional Kir2.1 under *GH146-GAL4* control caused early larval lethality, but this premature death could be circumvented with three tubulin promoter-driven copies of the temperature-sensitive repressor of GAL4, GAL80^ts^ (McGuire et al., 2003), which kept the expression of the channel at bay until the block was thermally relieved during adulthood.

Following their induction for 24 h at 31°C, both EGFP-tagged channels (Kir2.1 and Kir2.1-nc) were detected in PNs of 5 day-old adults at comparable levels and in the same anatomical distribution (Fig. 3*A*). Whole-cell current-clamp recordings showed that EGFP::Kir2.1 lowers the input resistance and membrane time constant relative to EGFP::Kir2.1-nc (Fig. 3*B*–*D*) and powerfully opposes depolarization: Kir2.1-expressing neurons required approximately two-fold larger depolarizing currents to drive spiking across a firing rate range of 1–50 Hz (Fig. 3*E*). Although Kir2.1 does not strictly silence the population of neurons in which it is expressed (the added potassium conductance can always be compensated by a large enough current injection; Figs. 3*B,E*), the currents necessary to do so seem difficult to attain *in vivo*.

**Figure 3.**
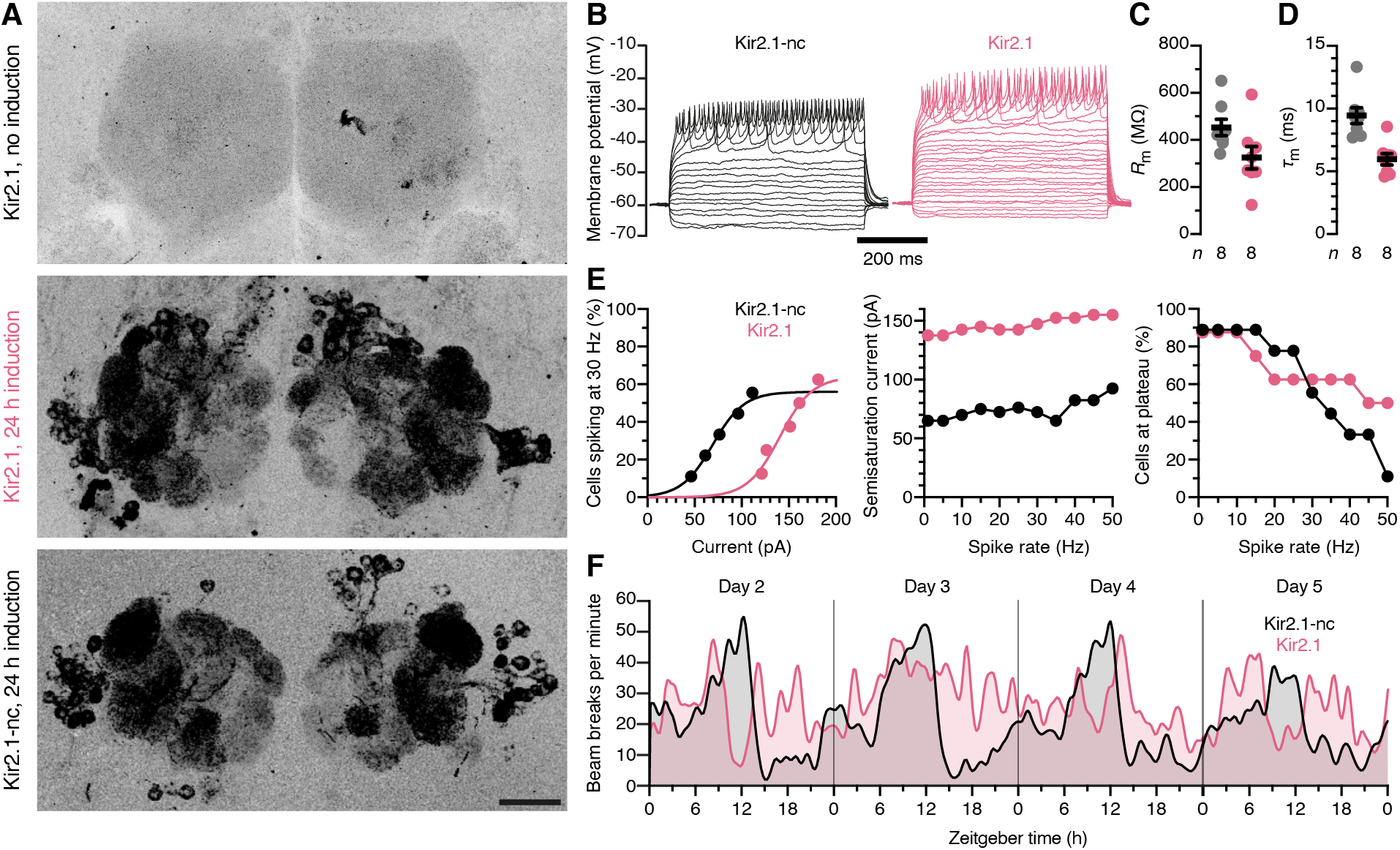
A transcriptomics-friendly neuronal activity clamp. ***A***, Maximum intensity projections of confocal image stacks through the antennal lobes of 5-day old female flies carrying *EGFP::Kir2.1* or *EGFP::Kir2.1-nc* transgenes under *GH146-GAL4* and *tub-GAL80^ts^* control. The expression of Kir2.1 constructs is undetectable at 21°C (top) but induced at 31°C (center and bottom). Scale bar, 20 μm. ***B***, Example voltage responses to 5-pA current steps of antennal lobe PNs expressing EGFP::Kir2.1-nc (black) or EGFP::Kir2.1 (red). ***C***, ***D***, Kir2.1 (red) lowers the input resistance *R*_m_ (*p* = 0.0481, *t*-test; ***C***) and shortens the membrane time constant *τ*_m_ (*p* = 0.0005, *t*-test; ***D***) relative to Kir2.1-nc (black). Circles, individual PNs; bars, means ± SEM. ***E***, Cumulative distribution functions of the percentages of PNs reaching a spike frequency of 30 Hz at different levels of injected current (left); semisaturation currents (center) and percentages of cells reaching spike rates of 1–50 Hz, for PNs expressing Kir2.1-nc (black) or Kir2.1 (red). ***F***, Circadian locomotor rhythms in constant darkness. Locomotion was quantified as the total number of midline crossings per minute in groups of 16 flies expressing Kir2.1-nc (black) or Kir2.1 (red) under *pdf-GAL4* control. The traces were smoothed with a Gaussian kernel (1.25 hours FWHM) and show data collected on days 2–5 after the flies were transferred to activity monitors.

A simple behavioral test supported this conclusion. Adult-onset expression of Kir2.1 in the PDF-expressing ventral subset of lateral pacemaker neurons (using the *pdf-GAL4* driver) disrupted the circadian locomotor rhythm in constant darkness, as expected (Nitabach et al., 2002), whereas flies expressing Kir2.1-nc remained rhythmic (Fig. 3*F*).

### Trans-synaptic regulation of gene expression: transmitter release and synapse remodeling, and a late shift to proteostasis and neuroprotection

To delineate changes in presynaptic gene expression after muting postsynaptic neural activity, we compared the third antennal segment transcriptomes of flies expressing either Kir2.1 or Kir2.1-nc in PNs (Fig. 4*A*). We studied three induction times—12 hours, 48 hours, and 96 hours—in individuals that were age-matched at the point of analysis: all tissues were harvested between ZT6 and ZT7 on the fifth post-eclosion day (Fig. 4*A*). Two sequencing technologies—3’ DGE for the 12-and 48-hour groups and standard RNA-seq for the 96-hour group—gave similar mapping metrics (Tables 3 and 4).

**Figure 4.**
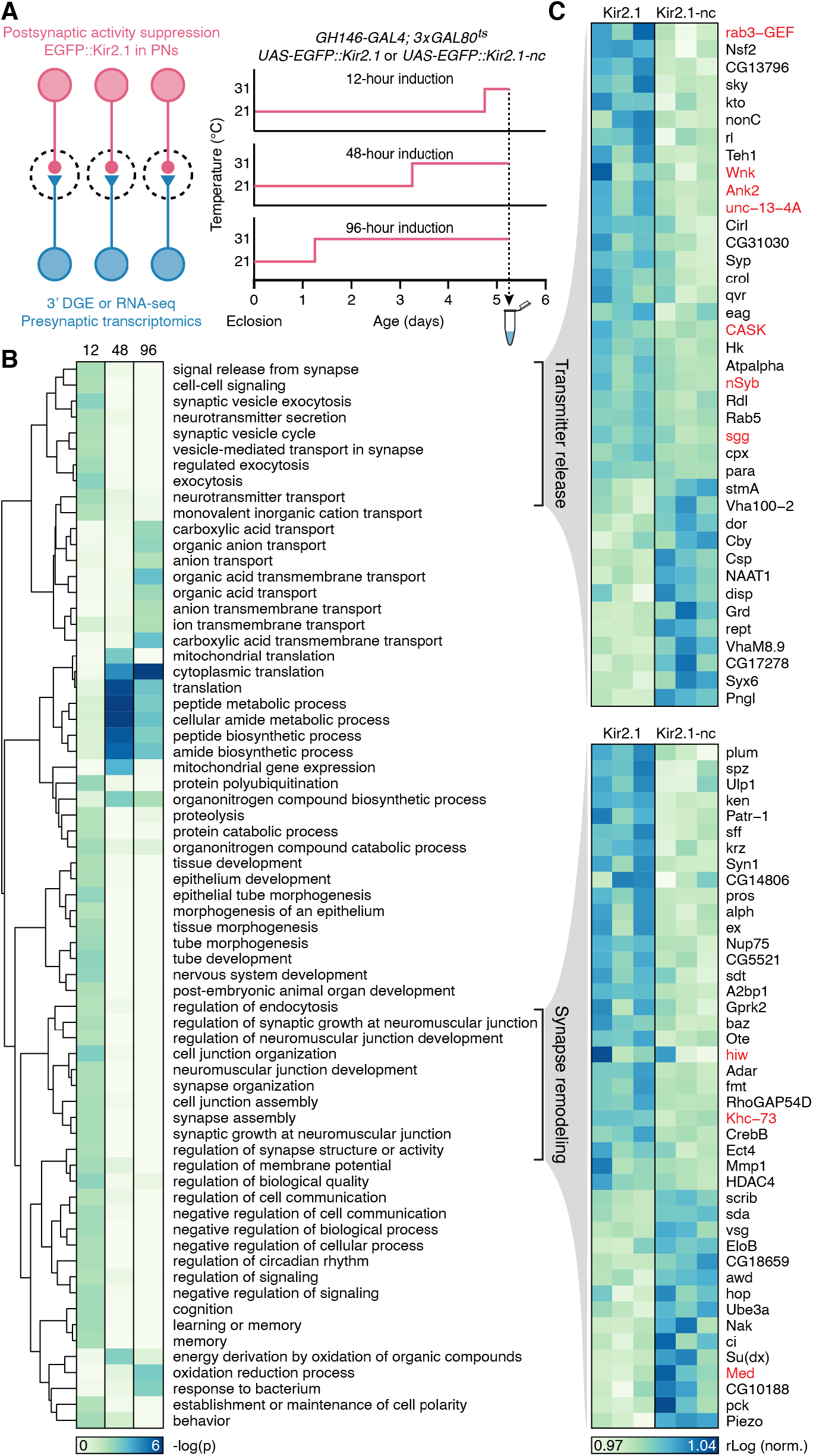
Trans-synaptic regulation of gene expression: transmitter release and synapse remodeling, and a late shift to proteostasis and neuroprotection. ***A***, Experimental design. **B.** Enrichment of GO biological process terms in third antennal segment transcriptomes after 12-hour, 48-hour, and 96-hour induction of Kir2.1. The dendrogram represents semantic groupings among GO terms. ***C***, Expression levels of transcripts attached to two semantic groupings, “transmitter release” and “synapse remodeling” (***B***), after 12 hours of induction of Kir2.1 or Kir2.1-nc. Each column represents a sequencing library. Gene products previously implicated in homeostatic synaptic plasticity (see Discussion) are indicated in red.

**Table 3.** Mapping metrics for third antennal segment transcriptomes collected after 12 or 48 hours of induction of Kir2.1 (K) or a non-conducting control (C)

| ID | *n* Reads | *n* Mapped | % Mapped | *n* Unique | % Unique | Average R<sup>2</sup> |
| --- | --- | --- | --- | --- | --- | --- |
| K12-1 | 29,660,291 | 15,855,218 | 53.46% | 14,082,433 | 47.48% | 0.9939 |
| K12-2 | 28,904,100 | 15,808,260 | 54.69% | 14,110,730 | 48.82% | 0.9943 |
| K12-3 | 28,938,152 | 15,549,498 | 53.73% | 13,323,889 | 46.04% | 0.9935 |
| C12-1 | 22,023,606 | 10,963,417 | 49.78% | 9,701,601 | 44.05% | 0.9941 |
| C12-2 | 15,850,790 | 8,864,492 | 55.93% | 7,922,966 | 49.99% | 0.9947 |
| C12-3 | 11,834,929 | 6,237,128 | 52.70% | 5,546,857 | 46.87% | 0.9946 |
| K48-1 | 28,004,316 | 18,815,975 | 67.19% | 16,487,747 | 58.88% | 0.9953 |
| K48-2 | 17,661,910 | 11,229,908 | 63.58% | 9,851,297 | 55.78% | 0.9951 |
| K48-3 | 27,158,740 | 15,643,356 | 57.60% | 13,588,111 | 50.03% | 0.9957 |
| C48-1 | 40,636,120 | 26,365,750 | 64.88% | 23,831,661 | 58.65% | 0.9770 |
| C48-2 | 26,993,657 | 13,344,752 | 49.44% | 11,474,648 | 42.51% | 0.9612 |
| C48-3 | 32,525,963 | 18,944,019 | 58.24% | 17,227,196 | 52.96% | 0.9787 |
| **Average** | **25,849,381** | **14,801,814** | **56.77%** | **13,095,761** | **50.17%** | **0.9890** |

Absolute number and percentages of reads aligned to the transcript file of the Berkeley Drosophila Genome Project (BDGP) release 5.74. *n* Reads: number of raw reads; *n* Mapped and % Mapped: number of reads mapping to the reference transcriptome with corresponding percentage; *n* Unique and % Unique: number of reads mapping to a unique location in the reference transcriptome with corresponding percentage; Average R<sup>2</sup>: average replicate correlation (Pearson's correlation coefficient calculated across all genes with a non-zero read count in at least one replicate).

**Table 4.** Mapping metrics for third antennal segment transcriptomes collected after 96 hours of induction of Kir2.1 (K) or a non-conducting control (C)

| ID | *n* Reads | *n* Mapped | % Mapped | *n* Unique | % Unique | Average R<sup>2</sup> |
| --- | --- | --- | --- | --- | --- | --- |
| K96-1 | 31,213,542 | 25,030,748 | 80.19% | 14,405,148 | 46.15% | 0.9836 |
| K96-2 | 55,190,242 | 43,522,921 | 78.86% | 26,736,203 | 48.44% | 0.9937 |
| K96-3 | 58,556,838 | 47,205,802 | 80.62% | 31,655,311 | 54.06% | 0.9941 |
| K96-4 | 29,803,652 | 24,632,464 | 82.65% | 17,888,586 | 60.02% | 0.9933 |
| K96-5 | 28,647,909 | 23,564,548 | 82.26% | 17,152,909 | 59.87% | 0.9937 |
| K96-6 | 31,030,362 | 23,420,613 | 75.48% | 17,208,103 | 55.46% | 0.9925 |
| K96-7 | 26,014,249 | 21,066,849 | 80.98% | 14,655,033 | 56.33% | 0.9932 |
| K96-8 | 30,622,108 | 24,934,568 | 81.43% | 17,368,348 | 56.72% | 0.9928 |
| K96-9 | 35,563,235 | 27,845,191 | 78.30% | 14,971,400 | 42.10% | 0.9869 |
| K96-10 | 26,388,734 | 21,106,014 | 79.98% | 13,725,816 | 52.01% | 0.9904 |
| C96-1 | 29,748,012 | 24,036,358 | 80.80% | 15,070,224 | 50.66% | 0.9928 |
| C96-2 | 61,127,179 | 48,487,739 | 79.32% | 30,257,167 | 49.50% | 0.9933 |
| C96-3 | 58,864,472 | 49,116,496 | 83.44% | 36,050,234 | 61.24% | 0.9939 |
| C96-4 | 30,582,966 | 25,278,565 | 82.66% | 17,865,165 | 58.42% | 0.9928 |
| C96-5 | 27,223,175 | 22,593,695 | 82.99% | 16,178,509 | 59.43% | 0.9935 |
| C96-6 | 29,951,768 | 24,706,089 | 82.49% | 17,520,016 | 58.49% | 0.9908 |
| C96-7 | 32,514,120 | 26,933,883 | 82.84% | 19,342,591 | 59.49% | 0.9940 |
| C96-8 | 27,902,225 | 23,365,715 | 83.74% | 17,388,108 | 62.32% | 0.9936 |
| C96-9 | 28,058,965 | 21,639,543 | 77.12% | 11,934,889 | 42.54% | 0.9911 |
| C96-10 | 28,737,234 | 23,114,269 | 80.43% | 14,720,484 | 51.22% | 0.9928 |
| **Average** | **35,387,049** | **28,580,104** | **80.83%** | **19,104,712** | **54.22%** | **0.9921** |

Absolute number and percentages of reads aligned to the transcript file of the Berkeley Drosophila Genome Project (BDGP) release 5.74. *n* Reads: number of raw reads; *n* Mapped and % Mapped: number of reads mapping to the reference transcriptome with corresponding percentage; *n* Unique and % Unique: number of reads mapping to a unique location in the reference transcriptome with corresponding percentage; Average R<sup>2</sup>: average replicate correlation (Pearson's correlation coefficient calculated across all genes with a non-zero read count in at least one replicate).

For 12-hour induction, experimental (Kir2.1) and control (Kir2.1-nc) flies were placed at 31°C from ZT18 until ZT6 on their fifth post-eclosion day and decapitated between ZT6 and ZT7 on the same day (Fig. 4*A*). Three biological replicates were sequenced for each genotype, one from males and two from females, and sex differences were accounted for and removed by the regression model entered into DESeq2. This analysis highlighted 25 differentially expressed antennal segment genes with FDR-adjusted *p* < 0.20 (Table 5). The average changes in absolute expression levels of the top 20 differentially expressed genes were about 15-fold smaller than those of the top 20 clock-controlled genes (log_2_ fold changes: 0.35 ± 0.08 for homeostatic genes vs.1.9 ± 0.9 for circadian genes; Tables 2 and 5), resulting in many fewer significant hits for the same FDR threshold. Among genes with the smallest FDR-adjusted *p*-values, many are involved in cell fate commitment and morphogenesis (Table 5); eight (*bazooka*, *sugar-free frosting*, *plum*, *prospero*, *Ankyrin 2*, *spätzle*, *Syncrip*, and *ATP6AP2*) have been linked to synaptic organization or synapse formation, however indirectly (Doe et al., 1991; Ruiz-Canada et al., 2004; Koch et al., 2008; Pielage et al., 2008; Baas et al., 2011; Sutcliffe et al., 2013; Yu et al., 2013; Halstead et al., 2014; Dubos et al., 2015).

**Table 5.** Differentially expressed genes after 12 hours of Kir2.1 induction (FDR-adjusted *p* < 0.20)

| Gene | Baseline expression | Log<sub>2</sub> fold change | $p$ | FDR | Description |
| --- | --- | --- | --- | --- | --- |
| *CG11550* | 7106.49 | -0.3861 | 3.07E-06 | 4.71E-03 | lipid binding |
| *Rbfox1* | 1799.04 | 0.3151 | 3.22E-05 | 1.35E-02 | RNA binding |
| *Trc8* | 885.38 | -0.5493 | 5.29E-05 | 1.35E-02 | protein ubiquitination |
| *baz* | 2163.42 | 0.3019 | 4.58E-05 | 1.35E-02 | cell polarity, junction formation |
| *Dso1* | 91551.91 | -0.2967 | 4.77E-05 | 1.35E-02 | immune response |
| *lncRNA: noe* | 89869.98 | 0.3187 | 1.95E-05 | 1.35E-02 | no functional information |
| *GstE3* | 2132.39 | 0.3211 | 2.19E-04 | 4.80E-02 | glutathione metabolism |
| *hth* | 11449.58 | 0.2245 | 3.91E-04 | 6.65E-02 | brain development |
| *sff* | 1232.13 | 0.4033 | 3.58E-04 | 6.65E-02 | NMJ development |
| *plum* | 981.88 | 0.4713 | 4.66E-04 | 7.14E-02 | axon pruning |
| *CG11873* | 977.85 | 0.3519 | 6.28E-04 | 8.74E-02 | stress response, transcription |
| *pros* | 1847.93 | 0.3498 | 9.49E-04 | 1.21E-01 | neuronal differentiation |
| *Ank2* | 4808.04 | 0.3167 | 1.68E-03 | 1.30E-01 | membrane scaffolding |
| *CG6421* | 1256.69 | -0.3262 | 1.70E-03 | 1.30E-01 | immune response |
| *Hcf* | 801.97 | 0.5215 | 1.44E-03 | 1.30E-01 | chromatic remodeling |
| *Kr-h2* | 805.79 | -0.3513 | 1.59E-03 | 1.30E-01 | membrane organization |
| *Pis* | 1370.97 | 0.2670 | 1.44E-03 | 1.30E-01 | signal transduction |
| *crol* | 1882.00 | 0.2505 | 1.69E-03 | 1.30E-01 | cell adhesion |
| *sdt* | 1635.14 | 0.3144 | 1.52E-03 | 1.30E-01 | cell polarity, junction formation |
| *spz* | 755.96 | 0.4628 | 1.34E-03 | 1.30E-01 | axon guidance, immune response |
| *Syp* | 3782.12 | 0.2909 | 1.84E-03 | 1.34E-01 | RNA binding, NMJ transmission |
| *ATP6AP2* | 1413.52 | -0.3679 | 2.08E-03 | 1.45E-01 | axonal transport |
| *CG9171* | 697.10 | 0.3263 | 2.27E-03 | 1.45E-01 | O-linked mannosylation |
| *RpL38* | 9770.50 | -0.2603 | 2.25E-03 | 1.45E-01 | translation |
| *mAcon1* | 1385.07 | -0.3415 | 2.80E-03 | 1.65E-01 | mitochondrial Krebs cycle |
| *CG3907* | 2306.39 | -0.2459 | 2.75E-03 | 1.65E-01 | no functional information |

$p$ , unadjusted $p$ -value; FDR, $p$ -value adjusted for multiple comparisons.

For 48-hour induction, experimental and control groups were shifted to 31°C at ZT6 of their third post-eclosion day and decapitated between ZT6 and ZT7 on day 5 (Fig. 4*A*). Three biological replicates—all from males—were sequenced for each genotype using 3’ DGE technology. One of the Kir2.1-nc replicates did not cluster well with the others (Table 3) and was excluded from the differential expression analysis, which produced 26 hits with FDR-adjusted *p* < 0.20 (Table 6). Conspicuous among these hits were several ribosomal components and three chaperones of the Hsp20 family (Hsp27, Hsp67Bc, and Hsp23) (Haslbeck et al., 2019). At first glance, the upregulation of heat shock proteins might suggest a direct effect of our method of transgene induction (31°C heat), but upon reflection heat cannot explain the observed differences because experimental and control flies were exposed to the same temperature regime. A more plausible explanation is, therefore, that prolonged postsynaptic silencing places an intense homeostatic burden on presynaptic partners which elicits a generalized increase in protein synthesis.

**Table 6.** Differentially expressed genes after 48 hours of Kir2.1 induction (FDR-adjusted *p* < 0.20)

| Gene | Baseline expression | Log<sub>2</sub> fold change | $p$ | FDR | Description |
| --- | --- | --- | --- | --- | --- |
| *CG8620* | 36.69 | 2.2870 | 1.36E-09 | 1.27E-05 | no functional information |
| *CG42502* | 7610.35 | 0.7250 | 7.72E-07 | 3.60E-03 | no functional information |
| *Hsp27* | 141.78 | 1.1780 | 8.78E-06 | 2.73E-02 | HSP20-like chaperone |
| *BomS3* | 287.22 | 1.0470 | 1.41E-05 | 3.28E-02 | immune response |
| *CR41619* | 337.93 | 0.9580 | 1.77E-05 | 3.30E-02 | no functional information |
| *Fbp2* | 49.24 | -1.3890 | 3.64E-05 | 5.65E-02 | alcohol dehydrogenase |
| *RpLP2* | 13612.76 | 0.7030 | 4.39E-05 | 5.84E-02 | structural constituent of ribosome |
| *Mst57Dc* | 10.73 | -1.5090 | 5.61E-05 | 5.91E-02 | mating behavior |
| *S-Lap4* | 8.45 | -1.4820 | 5.71E-05 | 5.91E-02 | proteolysis |
| *CG5986* | 566.50 | 1.0110 | 6.91E-05 | 6.44E-02 | RNA binding |
| *CG3224* | 171.53 | 0.9550 | 8.83E-05 | 7.48E-02 | ribosomal export |
| *CR31032* | 31.67 | 1.4380 | 1.04E-04 | 8.06E-02 | no functional information |
| *Trx-2* | 4120.04 | 0.6480 | 1.12E-04 | 8.06E-02 | thioredoxin, oxidative stress |
| *Cyp4g1* | 46.34 | -1.5000 | 1.23E-04 | 8.23E-02 | oxidation-reduction |
| *CG7920* | 1911.26 | -0.6730 | 1.59E-04 | 8.23E-02 | acetyl CoA metabolism |
| *Hsp67Bc* | 54.45 | 1.3110 | 1.51E-04 | 8.23E-02 | translation, protein folding |
| *Jon65Aiv* | 22.70 | -1.4610 | 1.52E-04 | 8.23E-02 | proteolysis |
| *mRpS5* | 722.02 | 0.7000 | 1.52E-04 | 8.23E-02 | structural constituent of ribosome |
| *RpS3A* | 31350.61 | 0.5610 | 1.81E-04 | 8.89E-02 | structural constituent of ribosome |
| *Tsp39D* | 1213.20 | 0.5520 | 2.08E-04 | 9.69E-02 | cell membrane scaffolding |
| *CG17454* | 4301.55 | 0.5740 | 3.44E-04 | 1.50E-01 | mRNA splicing |
| *CG6770* | 41760.18 | 0.4970 | 3.70E-04 | 1.50E-01 | transcriptional regulation |
| *Pkd2* | 10.79 | 1.3760 | 3.58E-04 | 1.50E-01 | cation transport |
| *Hsp23* | 141.98 | 1.0130 | 4.67E-04 | 1.81E-01 | protein folding |
| *Obp99d* | 176.07 | 1.0220 | 5.51E-04 | 1.98E-01 | smell perception |
| *snRNA:U2:34ABa* | 106.57 | 1.3060 | 5.49E-04 | 1.98E-01 | mRNA splicing |

$p$ , unadjusted $p$ -value; FDR, $p$ -value adjusted for multiple comparisons.

For 96-hour induction, experimental and control groups were kept at 31°C from ZT6 of their first post-eclosion day and again decapitated between ZT6 and ZT7 on day 5 (Fig. 4*A*). A total of 20 libraries were sequenced in two batches using RNA-seq technology. A different sequencing method was chosen to ensure that our results were valid across sequencing platforms, and more replicates were processed to increase sensitivity. The first batch consisted of 12 samples with six replicates from each of the two genotypes (all male third antennal segments). Two replicates of each genotype in the first batch (K96-2, K96-3, C96-2, and C96-3) were sequenced to twice the depth of the others to detect very lowly expressed genes more reliably. The second batch (eight samples in total) consisted of another four samples of each genotype, two each from females and two from males. Two samples (K96-1 and K96-9) had low within-batch correlations and were omitted from the analysis (Table 4). The increase in statistical power enabled the detection of 32 differentially expressed genes with FDR-adjusted *p* < 0.05 after controlling for sex and batch in DESeq2 (Table 7). Three biological processes stand out among these differentially expressed genes. First, six genes related to the Imd and Toll pathways of the innate immune response (Valanne et al., 2011) were strongly downregulated: the pattern recognition receptor PGRP-SD; the antibacterial peptide Drosocin (Dro); the negative regulator of Imd, pirk; and the antimicrobial peptides Bomanin Short 1, 3, and 5 (a.k.a. IM1, IM2, and IM3). Second, chaperones of the Hsp20 family, already encountered after 48-h induction, were again upregulated (Hsp26, Hsp23, and Hsp67Bc) (Haslbeck et al., 2019). And third, four genes involved in programmed cell death were differentially expressed, with two pro-apoptotic factors downregulated—matrix metalloproteinase 1 (Mmp 1) and apoptosis-inducing factor (AIF)—and two gene products inhibiting apoptosis up-regulated (Hsp26, Buffy) (Quinn et al., 2003; Wang et al., 2004; Joza et al., 2008). Overall, the 96-hour picture suggests a transcriptional landscape tilted toward cell protection and maintenance.

**Table 7.**
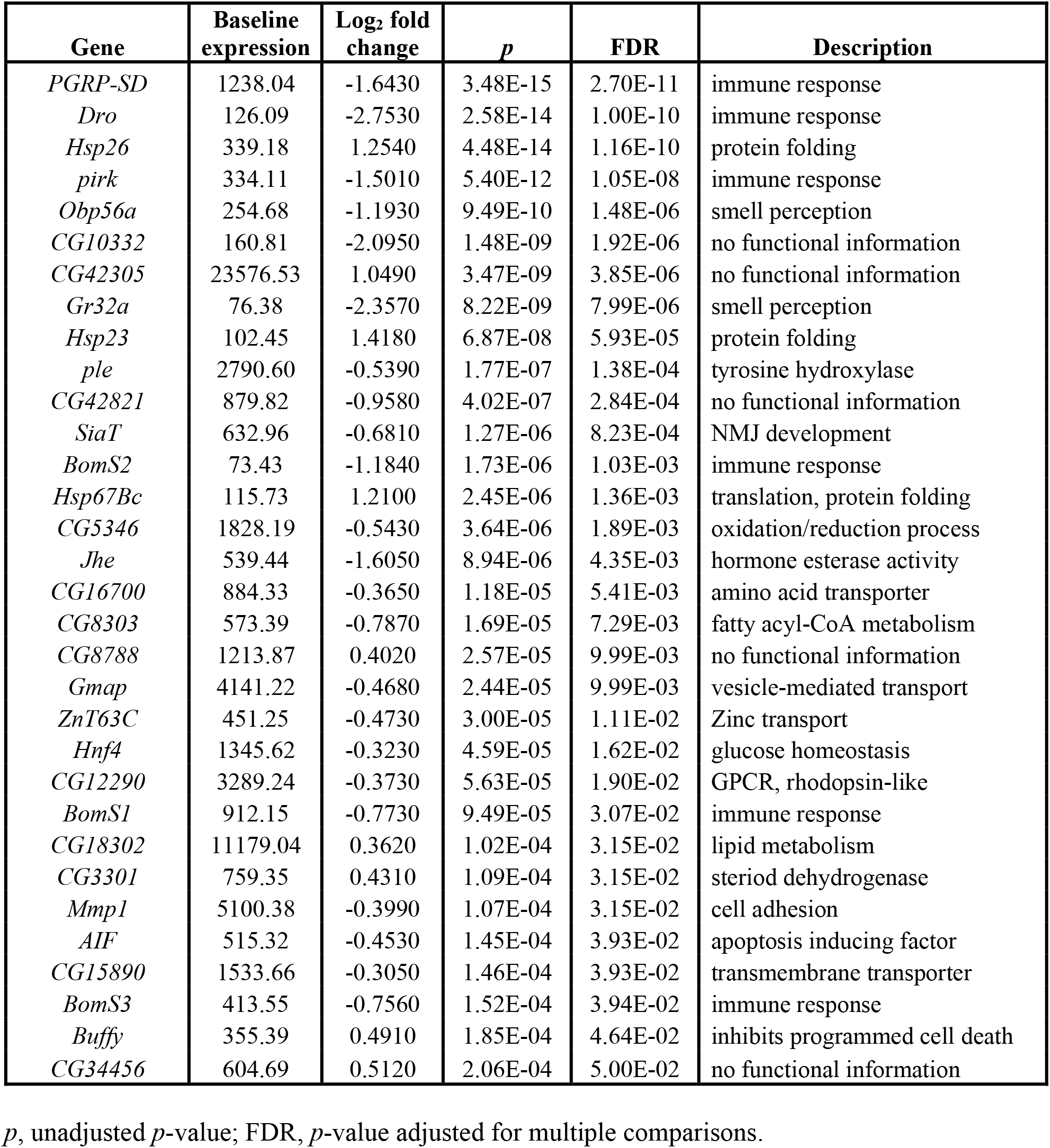
Differentially expressed genes after 96 hours of Kir2.1 induction (FDR-adjusted *p* < 0.05)

| Gene | Baseline expression | Log<sub>2</sub> fold change | $p$ | FDR | Description |
| --- | --- | --- | --- | --- | --- |
| *PGRP-SD* | 1238.04 | -1.6430 | 3.48E-15 | 2.70E-11 | immune response |
| *Dro* | 126.09 | -2.7530 | 2.58E-14 | 1.00E-10 | immune response |
| *Hsp26* | 339.18 | 1.2540 | 4.48E-14 | 1.16E-10 | protein folding |
| *pirk* | 334.11 | -1.5010 | 5.40E-12 | 1.05E-08 | immune response |
| *Obp56a* | 254.68 | -1.1930 | 9.49E-10 | 1.48E-06 | smell perception |
| *CG10332* | 160.81 | -2.0950 | 1.48E-09 | 1.92E-06 | no functional information |
| *CG42305* | 23576.53 | 1.0490 | 3.47E-09 | 3.85E-06 | no functional information |
| *Gr32a* | 76.38 | -2.3570 | 8.22E-09 | 7.99E-06 | smell perception |
| *Hsp23* | 102.45 | 1.4180 | 6.87E-08 | 5.93E-05 | protein folding |
| *ple* | 2790.60 | -0.5390 | 1.77E-07 | 1.38E-04 | tyrosine hydroxylase |
| *CG42821* | 879.82 | -0.9580 | 4.02E-07 | 2.84E-04 | no functional information |
| *SiaT* | 632.96 | -0.6810 | 1.27E-06 | 8.23E-04 | NMJ development |
| *BomS2* | 73.43 | -1.1840 | 1.73E-06 | 1.03E-03 | immune response |
| *Hsp67Bc* | 115.73 | 1.2100 | 2.45E-06 | 1.36E-03 | translation, protein folding |
| *CG5346* | 1828.19 | -0.5430 | 3.64E-06 | 1.89E-03 | oxidation/reduction process |
| *Jhe* | 539.44 | -1.6050 | 8.94E-06 | 4.35E-03 | hormone esterase activity |
| *CG16700* | 884.33 | -0.3650 | 1.18E-05 | 5.41E-03 | amino acid transporter |
| *CG8303* | 573.39 | -0.7870 | 1.69E-05 | 7.29E-03 | fatty acyl-CoA metabolism |
| *CG8788* | 1213.87 | 0.4020 | 2.57E-05 | 9.99E-03 | no functional information |
| *Gmap* | 4141.22 | -0.4680 | 2.44E-05 | 9.99E-03 | vesicle-mediated transport |
| *ZnT63C* | 451.25 | -0.4730 | 3.00E-05 | 1.11E-02 | Zinc transport |
| *Hnf4* | 1345.62 | -0.3230 | 4.59E-05 | 1.62E-02 | glucose homeostasis |
| *CG12290* | 3289.24 | -0.3730 | 5.63E-05 | 1.90E-02 | GPCR, rhodopsin-like |
| *BomS1* | 912.15 | -0.7730 | 9.49E-05 | 3.07E-02 | immune response |
| *CG18302* | 11179.04 | 0.3620 | 1.02E-04 | 3.15E-02 | lipid metabolism |
| *CG3301* | 759.35 | 0.4310 | 1.09E-04 | 3.15E-02 | steriod dehydrogenase |
| *Mmp1* | 5100.38 | -0.3990 | 1.07E-04 | 3.15E-02 | cell adhesion |
| *AIF* | 515.32 | -0.4530 | 1.45E-04 | 3.93E-02 | apoptosis inducing factor |
| *CG15890* | 1533.66 | -0.3050 | 1.46E-04 | 3.93E-02 | transmembrane transporter |
| *BomS3* | 413.55 | -0.7560 | 1.52E-04 | 3.94E-02 | immune response |
| *Buffy* | 355.39 | 0.4910 | 1.85E-04 | 4.64E-02 | inhibits programmed cell death |
| *CG34456* | 604.69 | 0.5120 | 2.06E-04 | 5.00E-02 | no functional information |

$p$ , unadjusted $p$ -value; FDR, $p$ -value adjusted for multiple comparisons.

To obtain an aerial overview of transcriptionally regulated biological processes during all induction periods, we probed for coordinated changes in functionally related sets of genes via gene ontology (GO) enrichment analyses. These analyses were performed on all differentially expressed genes with unadjusted *p* < 0.05 and included only GO terms with more than 40 attached genes; the enriched GO terms (Fisher’s exact test, *p* < 0.01) were then hierarchically clustered according to their semantic similarity (Wang et al., 2007; Brionne et al., 2019) (Fig. 4*B*). After 12 hours of induction, presynaptic transcriptional changes centered on genes encoding synaptic release and remodeling machinery; at 48 hours and beyond, protein synthesis and degradation, and energy metabolism, predominated (Fig. 4*B,C*, Tables 8–10). Closer scrutiny of the 81 genes responsible for the early enrichment of synaptic GO annotations (Fig. 4*C*, Tables 11 and 12) uncovered many with established roles in homeostatic plasticity at the NMJ (or with known interactions with such genes), as we discuss below. Although typical transcripts showed only modest expression level changes of 15–30%, their regulation was clearly visible across multiple libraries (Fig. 4*D*). This consistency across biological replicates, and the statistically verified overabundance of synaptic genes in the differentially expressed set with low unadjusted *p*-values (Tables 11 and 12), suggest a genuine signal.

**Table 8.** Enriched GO biological process terms after 12 hours of Kir2.1 induction.

| GO id | Term | Annotated | Observed | Expected | Fisher's *p* |
| --- | --- | --- | --- | --- | --- |
| GO:0034330 | cell junction organization | 273 | 27 | 14 | 1.20E-03 |
| GO:0006887 | exocytosis | 75 | 11 | 4 | 1.80E-03 |
| GO:0016079 | synaptic vesicle exocytosis | 44 | 8 | 2 | 1.90E-03 |
| GO:0035295 | tube development | 550 | 45 | 29 | 2.00E-03 |
| GO:0065008 | regulation of biological quality | 888 | 66 | 47 | 2.20E-03 |
| GO:0060562 | epithelial tube morphogenesis | 373 | 33 | 20 | 2.40E-03 |
| GO:0007399 | nervous system development | 843 | 63 | 45 | 2.50E-03 |
| GO:0000209 | protein polyubiquitination | 58 | 9 | 3 | 3.20E-03 |
| GO:0035239 | tube morphogenesis | 400 | 34 | 21 | 3.90E-03 |
| GO:0007611 | learning or memory | 119 | 14 | 6 | 4.00E-03 |
| GO:0050890 | cognition | 119 | 14 | 6 | 4.00E-03 |
| GO:1901565 | organonitrogen compound catabolic process | 327 | 29 | 17 | 4.20E-03 |
| GO:0006836 | neurotransmitter transport | 95 | 12 | 5 | 4.20E-03 |
| GO:0010648 | negative regulation of cell communication | 241 | 23 | 13 | 4.30E-03 |
| GO:0023057 | negative regulation of signaling | 241 | 23 | 13 | 4.30E-03 |
| GO:0045055 | regulated exocytosis | 50 | 8 | 3 | 4.40E-03 |
| GO:0048519 | negative regulation of biological process | 1047 | 74 | 56 | 4.40E-03 |
| GO:0042391 | regulation of membrane potential | 51 | 8 | 3 | 5.00E-03 |
| GO:0048729 | tissue morphogenesis | 483 | 39 | 26 | 5.00E-03 |
| GO:0007610 | behavior | 377 | 32 | 20 | 5.20E-03 |
| GO:0007269 | neurotransmitter secretion | 74 | 10 | 4 | 5.40E-03 |
| GO:0099643 | signal release from synapse | 74 | 10 | 4 | 5.40E-03 |
| GO:0034329 | cell junction assembly | 176 | 18 | 9 | 5.40E-03 |
| GO:0007613 | memory | 86 | 11 | 5 | 5.50E-03 |
| GO:0051124 | synaptic growth at NMJ | 111 | 13 | 6 | 5.70E-03 |
| GO:0007416 | synapse assembly | 137 | 15 | 7 | 5.80E-03 |
| GO:0007528 | neuromuscular junction development | 138 | 15 | 7 | 6.20E-03 |
| GO:0060429 | epithelium development | 730 | 54 | 39 | 6.40E-03 |
| GO:0008582 | regulation of synaptic growth at NMJ | 88 | 11 | 5 | 6.50E-03 |
| GO:0050808 | synapse organization | 221 | 21 | 12 | 6.60E-03 |
| GO:0050803 | regulation of synapse structure or activity | 139 | 15 | 7 | 6.60E-03 |
| GO:0048523 | negative regulation of cellular process | 914 | 65 | 49 | 6.80E-03 |
| GO:0042752 | regulation of circadian rhythm | 65 | 9 | 3 | 6.90E-03 |
| GO:0048569 | post-embryonic animal organ development | 371 | 31 | 20 | 7.40E-03 |
| GO:0007267 | cell-cell signaling | 326 | 28 | 17 | 7.50E-03 |
| GO:0009888 | tissue development | 786 | 57 | 42 | 7.70E-03 |
| GO:0099003 | vesicle-mediated transport in synapse | 78 | 10 | 4 | 7.80E-03 |
| GO:0099504 | synaptic vesicle cycle | 78 | 10 | 4 | 7.80E-03 |
| GO:0010646 | regulation of cell communication | 595 | 45 | 32 | 8.70E-03 |
| GO:0023051 | regulation of signaling | 595 | 45 | 32 | 8.70E-03 |
| GO:0007163 | establishment or maintenance of cell polarity | 171 | 17 | 9 | 9.00E-03 |
| GO:0015672 | monovalent inorganic cation transport | 92 | 11 | 5 | 9.10E-03 |
| GO:0030163 | protein catabolic process | 228 | 21 | 12 | 9.30E-03 |
| GO:0002009 | morphogenesis of an epithelium | 470 | 37 | 25 | 9.30E-03 |
| GO:0006508 | proteolysis | 471 | 37 | 25 | 9.60E-03 |
| GO:1904396 | regulation of NMJ development | 93 | 11 | 5 | 9.80E-03 |
| GO:0030100 | regulation of endocytosis | 46 | 7 | 2 | 1.00E-02 |

Annotated, number of genes in the universe attached to a GO term; Observed, number of differentially expressed genes (unadjusted $p < 0.05$ ) attached to a GO term; Expected, number of genes expected by chance to be attached to a GO term

**Table 9.** Enriched GO biological process terms after 48 hours of Kir2.1 induction.

| GO id | Term | Annotated | Observed | Expected | Fisher's *p* |
| --- | --- | --- | --- | --- | --- |
| GO:0043603 | cellular amide metabolic process | 405 | 53 | 25 | 8.70E-08 |
| GO:0006518 | peptide metabolic process | 355 | 46 | 22 | 9.00E-07 |
| GO:0006412 | translation | 265 | 37 | 16 | 1.90E-06 |
| GO:0043043 | peptide biosynthetic process | 293 | 39 | 18 | 3.40E-06 |
| GO:0043604 | amide biosynthetic process | 311 | 40 | 19 | 5.90E-06 |
| GO:0002181 | cytoplasmic translation | 100 | 18 | 6 | 3.30E-05 |
| GO:0140053 | mitochondrial gene expression | 86 | 15 | 5 | 2.10E-04 |
| GO:0032543 | mitochondrial translation | 79 | 13 | 5 | 9.90E-04 |
| GO:1901566 | organonitrogen compound biosynthetic process | 554 | 52 | 34 | 1.28E-03 |
| GO:0015980 | energy derivation by oxidation of organic compounds | 54 | 10 | 3 | 1.50E-03 |

Annotated, number of genes in the universe attached to a GO term; Observed, number of differentially expressed genes (unadjusted $p < 0.05$ ) attached to a GO term; Expected, number of genes expected by chance to be attached to a GO term

**Table 10.** Enriched GO biological process terms after 96 hours of Kir2.1 induction.

| GO id | Term | Annotated | Observed | Expected | Fisher's *p* |
| --- | --- | --- | --- | --- | --- |
| GO:0002181 | cytoplasmic translation | 100 | 28 | 8 | 5.50E-09 |
| GO:1903825 | organic acid transmembrane transport | 42 | 11 | 4 | 4.90E-04 |
| GO:1905039 | carboxylic acid transmembrane transport | 42 | 11 | 4 | 4.90E-04 |
| GO:0043603 | cellular amide metabolic process | 405 | 53 | 34 | 5.70E-04 |
| GO:0043604 | amide biosynthetic process | 311 | 43 | 26 | 6.10E-04 |
| GO:0006412 | translation | 265 | 38 | 22 | 6.30E-04 |
| GO:0006518 | peptide metabolic process | 355 | 47 | 30 | 9.10E-04 |
| GO:0055114 | oxidation-reduction process | 308 | 42 | 26 | 9.40E-04 |
| GO:0043043 | peptide biosynthetic process | 293 | 40 | 25 | 1.22E-03 |
| GO:0009617 | response to bacterium | 158 | 25 | 13 | 1.32E-03 |
| GO:0015711 | organic anion transport | 89 | 16 | 7 | 2.59E-03 |
| GO:0046942 | carboxylic acid transport | 59 | 12 | 5 | 3.03E-03 |
| GO:0015849 | organic acid transport | 60 | 12 | 5 | 3.51E-03 |
| GO:1901566 | organonitrogen compound biosynthetic process | 554 | 63 | 46 | 6.21E-03 |
| GO:0034220 | ion transmembrane transport | 160 | 23 | 13 | 6.94E-03 |
| GO:0098656 | anion transmembrane transport | 58 | 11 | 5 | 7.74E-03 |
| GO:0006820 | anion transport | 118 | 18 | 10 | 8.80E-03 |

Annotated, number of genes in the universe attached to a GO term; Observed, number of differentially expressed genes (unadjusted $p < 0.05$ ) attached to a GO term; Expected, number of genes expected by chance to be attached to a GO term

**Table 11.** Differentially expressed genes (*p* < 0.05) attached to GO biological process terms in the semantic grouping “transmitter release” after 12 hours of Kir2.1 induction.

| Gene | Baseline expression | Log<sub>2</sub> fold change | $p$ |
| --- | --- | --- | --- |
| *Ank2* | 4808.04 | 0.3167 | 1.68E-03 |
| *crol* | 1882.00 | 0.2505 | 1.69E-03 |
| *Syp* | 3782.12 | 0.2909 | 1.84E-03 |
| *ATP6AP2* | 1413.52 | -0.3679 | 2.08E-03 |
| *Pngl* | 165.57 | -0.5711 | 2.84E-03 |
| *Nsf2* | 57.11 | 1.0758 | 4.99E-03 |
| *sky* | 574.22 | 0.4143 | 5.49E-03 |
| *CG13796* | 98.20 | 0.7603 | 5.64E-03 |
| *rab3-GEF* | 74.57 | 0.9719 | 5.81E-03 |
| *Atpalpha* | 25649.53 | 0.2121 | 7.80E-03 |
| *rept* | 43.03 | -0.9271 | 1.06E-02 |
| *nSyb* | 7467.28 | 0.1791 | 1.25E-02 |
| *Cirl* | 753.00 | 0.2911 | 1.37E-02 |
| *Teh1* | 490.53 | 0.3833 | 1.47E-02 |
| *CG17278* | 103.00 | -0.6499 | 1.91E-02 |
| *Grd* | 36.96 | -0.9307 | 1.93E-02 |
| *Cby* | 442.89 | -0.3202 | 2.13E-02 |
| *Syx6* | 229.68 | -0.4827 | 2.22E-02 |
| *rl* | 129.67 | 0.5579 | 2.28E-02 |
| *Vha100-2* | 1861.65 | -0.2887 | 2.39E-02 |
| *eag* | 1418.20 | 0.2192 | 2.62E-02 |
| *NAAT1* | 301.63 | -0.3582 | 2.92E-02 |
| *qvr* | 751.52 | 0.2494 | 2.93E-02 |
| *Rab5* | 6457.41 | 0.1526 | 3.09E-02 |
| *kto* | 125.71 | 0.5835 | 3.33E-02 |
| *Csp* | 246.20 | -0.3718 | 3.50E-02 |
| *Rdl* | 9593.74 | 0.1719 | 3.67E-02 |
| *para* | 15702.82 | 0.1020 | 3.70E-02 |
| *CASK* | 2105.66 | 0.1889 | 3.73E-02 |
| *stmA* | 1105.17 | -0.2661 | 4.04E-02 |
| *Hk* | 1229.40 | 0.1803 | 4.22E-02 |
| *disp* | 51.74 | -0.7734 | 4.23E-02 |
| *nonC* | 55.67 | 0.7504 | 4.33E-02 |
| *unc-13-4A* | 244.53 | 0.3676 | 4.53E-02 |
| *Wnk* | 233.48 | 0.4181 | 4.57E-02 |
| *cpx* | 9807.02 | 0.1367 | 4.78E-02 |
| *sgg* | 3797.38 | 0.1451 | 4.83E-02 |
| *CG31030* | 278.71 | 0.3348 | 4.92E-02 |
| *dor* | 43.31 | -0.7143 | 4.99E-02 |

**Table 12.**
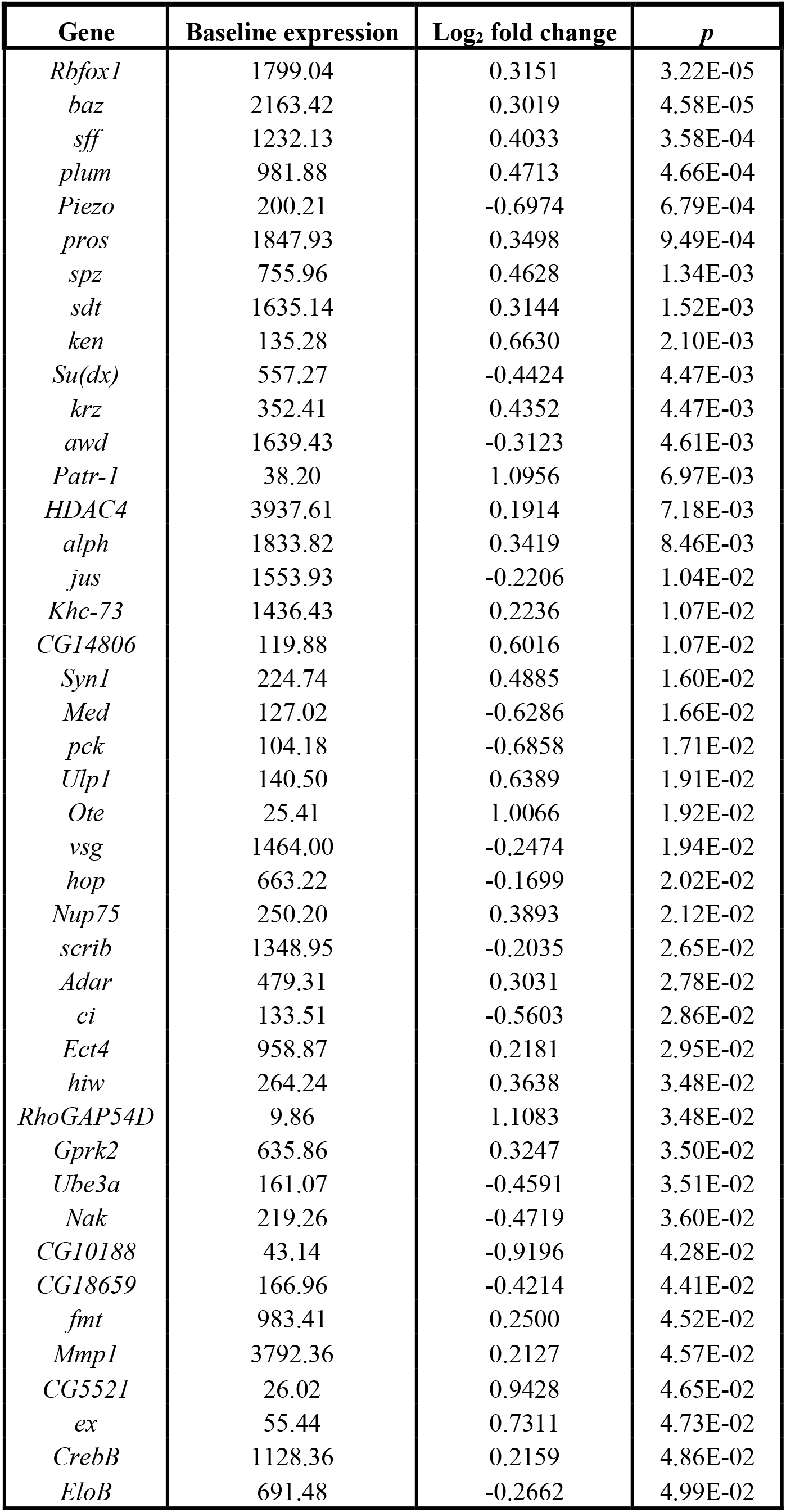
Differentially expressed genes (*p* < 0.05) attached to GO biological process terms in the semantic grouping “synapse remodeling” after 12 hours of Kir2.1 induction.

### Cross-validation of regulated genes with 3’ DGE, RNA-seq, and RT-qPCR

As a further validation of our gene expression measurements, we compared transcriptome-wide 3’ DGE with transcriptome-wide RNA-seq data. There was an approximately linear relationship between the average expression levels of all genes in all samples (Fig. 5*A*), with a small departure in lowly expressed genes caused by the extra amplification step in the RNA-seq protocol (see Methods); as a result RNA-seq reported systematically higher expression levels for scarce transcripts than did 3’ DGE. For genes transcribed at moderate to high expression levels, the two sequencing platforms were in close agreement.

**Figure 5.**
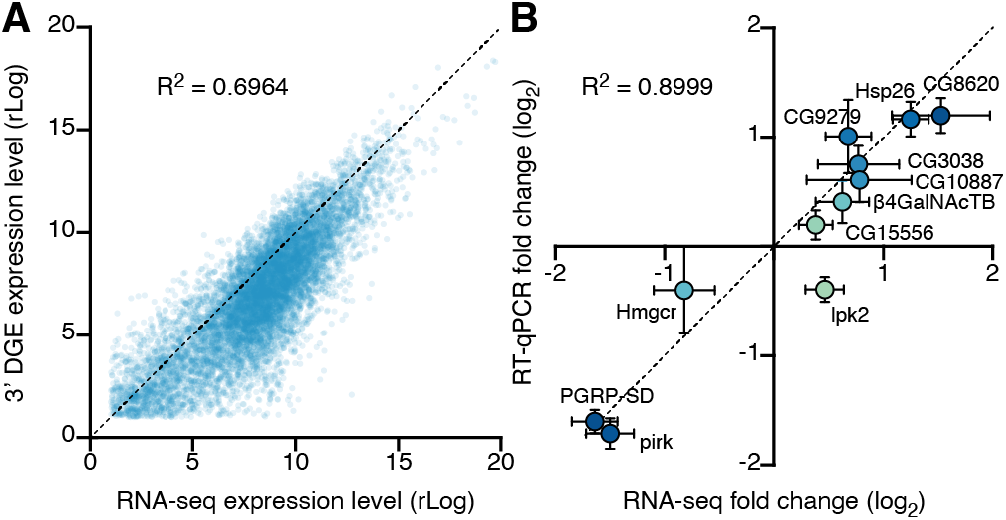
Cross-validation of differential gene expression. ***A***, Scatterplot of average gene expression levels determined by RNA-seq vs. 3’ DGE. ***B***, Scatterplot of log_2_ fold changes (means ± SEM) in the expression levels of 11 transcripts after 96-hour induction of Kir2.1, determined by RNA-seq or RT-qPCR.

We next selected 11 transcripts for RT-qPCR verification. These transcripts were chosen from the set of differentially expressed genes (unadjusted *p* < 0.05; both up- and down-regulated) in the 96-hour induction group and used to validate all deep sequencing data. The fold changes of the 11 chosen transcripts, as estimated by RT-qPCR with normalization to the housekeeping gene *CycK*, correlated tightly with 3’ DGE and RNA-seq measurements (Fig. 5*B*). This agreement between three independent measures of gene expression, at a transcriptome-wide scale and across several individual genes, lends confidence to our analysis.

## Discussion

### Trans-synaptic regulation of gene expression

Our study introduces an experimental system for detecting changes in gene expression in response to changes in the electrical excitability of a partner cell. The product of the *Kir2.1* transgene powerfully suppresses the activity of neurons in which it is expressed, while a control transgene, which codes for a potassium channel with a single amino acid substitution in its selectivity filter (Kir2.1-nc), has no effect (Fig. 3*B*–*F*). Isogenic strains expressing one or the other of these transgenes from the same chromosomal locus offer an ideal platform for differential gene expression analyses because differences between them can be pinned to a single codon change in the genome. The finding that prolonged postsynaptic silencing induces the expression of Hsp20 proteins in a manner unrelated to heat shock (Tables 6 and 7) underscores the power of this carefully controlled system.

The same finding, however, also highlights a limitation particular to our current approach. We imposed the Kir2.1 clamp on the first synaptic relay in the *Drosophila* olfactory system because its pre- and postsynaptic elements are easily separable by purely physical means, but this convenience exacted a price: the third antennal segment contains not only ORNs but also glial and support cells, which account for about two thirds of the segment’s cell population (Vosshall et al., 1999). We are therefore unable to determine whether the expression of Hsp20 proteins is exclusively or even partially neuronal. Although the same reservation does not apply to the many synaptic genes that are differentially expressed during the early phase of the homeostatic response (Fig. 4*B,C*), the presence of non-neuronal contaminants may nevertheless have hindered the detection of low-abundance neuronal transcripts or underestimated their fold change. Both of these drawbacks could be overcome by FACS-isolation of a genetically labeled cell population before RNA extraction, as would be required as a matter of course in all instances where the synaptic partners are anatomically intermingled. With this extra step, our system will be easily adapted for analyses of transcriptional changes elicited in presynaptic cells by a loss of postsynaptic responsivity, in postsynaptic cells by a loss of presynaptic input, or in glial cells by a heightened demand for synaptic remodeling.

Despite these caveats, many of the early expression level changes we detect affect genes encoding synaptic proteins with known, suspected, or at least plausible roles in homeostatic plasticity (Fig. 4*B,C*, Tables 5, 11, and 12) (Davis and Müller, 2015): elements of the wingless signaling system (e.g., Wnk, sgg), which acts as an endogenous suppressor of homeostatic compensation at the NMJ (Marie et al., 2010); the v-SNARE synaptobrevin (nSyb) and its chaperone Nsf2 (Söllner et al., 1993; Bacci et al., 2001); rab3 guanine nucleotide exchange factor (rab3-GEF), which controls the assembly and distribution of active zone components (Bae et al., 2016) and regulates the nucleotide state-dependent association of rab3 with synaptic vesicles, which in turn determines the calcium sensitivity of their release (Geppert et al., 1997; Müller et al., 2011); an active zone resident (unc-13-4a) known to associate with the Rab3-interacting molecule RIM and other active zone components (Schoch et al., 2002; Liu et al., 2011; Müller et al., 2012); a kinesin motor heavy chain (Khc-73) implicated in active zone assembly and synaptic homeostasis (Tsurudome et al., 2010); an active zone-integral guanylate kinase (CASK) that serves as a phosphorylation target of CDK5 (Samuels et al., 2007), which homeostatically regulates presynaptic calcium influx and release probability (Seeburg et al., 2008; Kim and Ryan, 2010); the E3 ubiquitin-protein ligase highwire (hiw) and the Smad protein Medea (Med), which in motor neuron terminals are part of the transduction cascade for a retrograde signal from muscle (Haghighi et al., 2003; McCabe et al., 2004; Goold and Davis, 2007); the cytoskeletal anchor Ankyrin 2 (Koch et al., 2008; Pielage et al., 2008); and subunits or accessory proteins of voltage-gated ion channels (quiver, ether-á-go-go, Hyperkinetic, paralytic) (Tables 5, 11, and 12). Collectively, these changes could signal an increase in the number of release sites or an expansion of the release-ready vesicle pool, inferred to represent the dominant quantal parameter change during homeostatic matching at ORN-to-PN synapses (Kazama and Wilson, 2008) and one of two homeostatic levers at the NMJ (the other being modulation of calcium influx into the terminal) (Müller et al., 2012).

When drawing comparisons with earlier work, however, it is important to bear in mind experimental differences in the speed of induction and expression of the homeostatic response. Abrupt adult-onset PN silencing resembles an acute postsynaptic receptor blockade at the NMJ more closely than it does the slow developmental processes studied in analyses of arbor size matching in the antennal lobe (Kazama and Wilson, 2008; Mosca and Luo, 2014), but homeostatic compensation at the NMJ is evident within minutes, long before changes in gene expression can occur (Frank et al., 2006). That elements of the homeostatic machinery are encoded by trans-synaptically regulated genes must therefore reflect a secondary layer of feedback control or a more profound reallocation of ORN synapses between PNs and other postsynaptic partners, such as local neurons of the antennal lobe (Groschner and Miesenböck, 2019).

Because changes in the expression levels of putative homeostatic genes are small compared to those of circadian-regulated genes (Figs. 1*D*, 4*C*), we were forced to apply lenient FDR thresholds to the 12-hour and 48-hour induction experiments, raising the specter of false positives in these data sets. Two observations should allay this concern. First, the 96-hour induction experiment, whose greater statistical power made the application of a more stringent significance threshold possible, recovered many of the same biological processes and indeed the same genes (e.g., Hsp23, Hsp67Bc) as the statistically weaker 48-hour induction experiment (Tables 9 and 10). Second, our RT-qPCR validation included several genes that failed to cross the most stringent FDR threshold (Fig. 5*B*). These RT-qPCR spot checks confirmed that expression level changes detected by RNA-seq or 3’ DGE were accurate. Nonetheless, new candidates emerging from our screen will need to survive rigorous functional studies before joining the ranks of established homeostatic plasticity genes.

### Labeling connections with trans-synaptically regulated genes?

Transcriptional changes that are controlled by trans-synaptic signals could be exploited for the generation of new circuit-breaking tools. In most neurobiological studies, the object of interest is not a population of genetically homogeneous neurons but an operational unit—a circuit—defined by connectivity rather than a common genetic marker (Miesenböck and Kevrekidis, 2005). Circuit analyses have benefited greatly from the development of trans-synaptic vectors (Card et al., 1990; Kuypers and Ugolini, 1990; Strack and Loewy, 1990; Wickersham et al., 2007), which travel along synaptic connections between specific types of neuron and serve as vehicles for the distribution of other encodable tools (Sjulson et al., 2016).

Ideally, trans-synaptic expression systems possess a mechanism that allows their initialization at a specific location, a rule that governs their propagation in the network, and gain. Viruses have some of these characteristics (Card et al., 1990; Kuypers and Ugolini, 1990; Strack and Loewy, 1990; Wickersham et al., 2007). Their infectious spread can follow routes of synaptic transmission, and replicative gain (where permitted) allows each infected neuron to supply more viral particles to its outputs than it receives from its inputs. Viral infections are, however, difficult to control and initialize with single-cell resolution and can produce considerable toxicity and extrasynaptic spread.

Lectins and catalytically crippled neurotoxins are also ferried across synapses (Schwab et al., 1979; Gerfen et al., 1982; Ruda and Coulter, 1982). These stripped-down trans-synaptic tracers lack the cytotoxic effects of viral replication but also the associated gain and the capacity to serve as gene delivery vehicles. The need to carry the label as payload across the synaptic cleft requires high expression levels, which jeopardize the specificity of transfer. Clearly, the ideal trans-synaptic vector would, instead of carrying its own genetic material or marker, act on expression cassettes that lie dormant in the genome of the host organism until switched on by a trans-synaptic signal.

Circuit-tracing systems such as *trans-*Tango, TRACT, and BAcTrace are built on this principle but require the reconstitution of an exogenous, contact- or ligand-based, cell-to-cell signaling apparatus (Huang et al., 2017; Talay et al., 2017; Cachero et al., 2020). This introduces additional genetic complexity and the danger of overexpression artefacts if the foreign molecules escape synaptic confinement. Eavesdropping on endogenous trans-synaptic communication during homeostatic plasticity offers a possible cure for these problems. Imagine a sudden, targeted loss of excitability in a small group of neurons or even a single cell, brought about by the inducible expression of Kir2.1. If presynaptic partners sense this perturbation and compensate homeostatically, the upregulation of plasticity genes could be coupled to the expression of sensors, actuators, transcription factors, or recombinases (Sjulson et al., 2016).

The chief obstacle to the development of this retrograde tracing technology is the small, at most two-fold, changes in homeostatic gene expression we detect (Tables 5, 11, and 12). We suspect that these changes would need to be amplified with adequate signal-to-noise ratio, perhaps by flipping a permanent recombination switch (Sjulson et al., 2016), to be practically useful. Region- or cell-specific differences in the capacity or mechanisms of homeostatic compensation are another potential concern. For example, it is likely that different plasticity mechanisms operate at excitatory and inhibitory synapses (or that the same genes respond to activity perturbations in opposite directions) (Turrigiano, 2011), leaving our hypothetical tracing tool blind to—or, depending on perspective, selective for—one or the other class of input. Still, the substantial overlap between components of the homeostatic machinery at the NMJ (Davis and Müller, 2015) and homeostatically regulated genes in the antennal lobe (Fig. 4*C*) suggests at least a measure of functional conservation from peripheral to central synapses. And, the activity-dependent changes in the expression levels of immediate early genes, which are widely used to capture task-specific neural ensembles (Morgan and Curran, 1991; Sjulson et al., 2016; DeNardo and Luo, 2017), are roughly equal to those of the trans-synaptically regulated genes we have identified.

## Acknowledgments

This work was supported by grants from the Wellcome Trust (106988/Z/15/Z, 090309/Z/09/Z, 089270/Z/09/Z) and the Gatsby Charitable Foundation (GAT3237). Paul Overton provided advice on RNA isolation; Amélie Baud gave many helpful tips for the gene ontology analysis; Ruth Brain assisted with stock maintenance and dissections; and Jessica Beevers helped with dissections and made delicious fly food (from the flies’ perspective).

## Author contributions

E.R.H. and G.M. designed the study, analyzed and interpreted the results, and wrote the paper. E.R.H. performed all experiments with the exception of electrophysiological recordings, which were done by D.P.

